# Pyrimidine biosynthesis inhibitors synergize with nucleoside analogs to block SARS-CoV-2 infection

**DOI:** 10.1101/2021.06.24.449811

**Authors:** David C. Schultz, Robert M. Johnson, Kasirajan Ayyanathan, Jesse Miller, Kanupriya Whig, Brinda Kamalia, Mark Dittmar, Stuart Weston, Holly L. Hammond, Carly Dillen, Lauren Castellana, Jae Seung Lee, Minghua Li, Emily Lee, Samuel Constant, Marc Ferrer, Christoph A. Thaiss, Matthew B. Frieman, Sara Cherry

## Abstract

The ongoing COVID-19 pandemic has highlighted the dearth of approved drugs to treat viral infections, with only ∼90 FDA approved drugs against human viral pathogens. To identify drugs that can block SARS-CoV-2 replication, extensive drug screening to repurpose approved drugs is underway. Here, we screened ∼18,000 drugs for antiviral activity using live virus infection in human respiratory cells. Dose-response studies validate 122 drugs with antiviral activity and selectivity against SARS-CoV-2. Amongst these drug candidates are 16 nucleoside analogs, the largest category of clinically used antivirals. This included the antiviral Remdesivir approved for use in COVID-19, and the nucleoside Molnupirivir, which is undergoing clinical trials. RNA viruses rely on a high supply of nucleoside triphosphates from the host to efficiently replicate, and we identified a panel of host nucleoside biosynthesis inhibitors as antiviral, and we found that combining pyrimidine biosynthesis inhibitors with antiviral nucleoside analogs synergistically inhibits SARS-CoV-2 infection in vitro and in vivo suggesting a clinical path forward.

## Main Text

The SARS-CoV-2 virus has infected more than 165 million people and led to more than 3 million deaths in the last year (*1*) (WHO.org). SARS-CoV-2-infected individuals typically develop mild to severe flulike symptoms, while a subset of individuals including the elderly or those harboring underlying chronic health complications, such as diabetes and heart disease, are particularly prone to develop severe to fatal clinical outcomes (*2*). While vaccines have been rapidly developed to combat SARS-CoV-2, there has been a dearth of antiviral therapeutics that have shown efficacy. There is an urgent need for therapeutics against SARS-CoV-2 which has been amplified by the emerging threats of variants that may evade vaccines. Large scale efforts are underway to identify antiviral drugs.

SARS-CoV-2 is a coronavirus, which is a family of single stranded positive sense RNA viruses, at least seven of which infect humans. RNA viruses including coronaviruses replicate using a virally encoded RNA-dependent RNA polymerase (RdRp), and nucleoside analogs which are misincorporated by the RdRp into the growing viral RNA chain are a large class of approved direct acting antivirals (*3*). Depending on the analog, misincorporation can lead to chain termination or mutagenesis ultimately inhibiting viral replication (*4*). Importantly, RdRps have conserved structures, and thus nucleoside analogs can show broad activity across related and unrelated viruses (*5, 6*). Therefore, repurposing efforts have identified nucleoside analogs that are active against newly emerging viruses and such efforts discovered that the nucleoside analog Remdesivir (Veklury^™^) inhibits SARS-CoV-2 replication becoming the first approved antiviral therapeutic against this novel coronavirus (*7, 8*).

Since all viruses, including SARS-CoV-2, are dependent on diverse cellular factors and metabolic products for their replication, the identification of host directed antivirals also shows promise. In particular, host nucleoside biogenesis is required for viral replication as RNA viruses require high levels of nucleoside triphosphates (NTPs) for their growth. Widespread efforts are underway to identify essential host pathways that are druggable, and to repurpose therapeutics against these host targets. Respiratory epithelial cells are the major cellular target for SARS-CoV-2 *in vivo*. We and others have found that the cellular entry pathway as well as other host-dependent steps in the viral lifecycle show cell-type specific differences and we were most interested in those antivirals that would be active in the respiratory tract (*9*). Therefore, we utilized the human respiratory cell line Calu-3 for these studies We set out to screen small molecule libraries that contained approved drugs, drugs in clinical trials and drugs with known targets to uncover both direct-acting and host directed antivirals using wild type virus and a cell-based, high-content assay in respiratory cells. We optimized a microscopy-based assay to achieve robust screening parameters (Z’>0.5) using vehicle (DMSO) and Remdesivir (10uM) as controls on each plate. We infect Calu-3 cells pre-treated with compounds and quantify infection of wild type SARS-CoV-2 48h post infection (hpi) using an antibody to dsRNA, a viral replication intermediate (*9*). In addition, we quantified the number of cells in each well to remove drugs that are cytotoxic.

We screened ∼18,000 drugs from three repurposing libraries: an in-house PENN library of ∼3,500 drugs, ∼3,400 drugs from the NCATS repurposing collection chosen to avoid overlap with the PENN library, and the ReFrame collection of ∼11,300 drugs most of which have been tested in human (*10*). Altogether, we have screened a large fraction of the drugs that can potentially be rapidly repurposed in respiratory cells with live virus complementing previous screening efforts (*9, 11,12*). We screened the PENN library in duplicate at 0.8 µM and identified 77 drugs that inhibit infection >60% and that had little toxicity (>80% cell number) in each screen; the NCATS library at 0.8 µM and identified 45 drugs that inhibit infection >60% and had little toxicity (>80% cell number); and the ReFrame library in duplicate at 3.6 µM and identified 135 compounds that inhibit infection >60% and had low toxicitity (>60% cell number) in both replicates (Fig S1a-c). We validated the candidates by repurchasing powders (PENN, NCATS) or testing pre-spotted validation plates (ReFrame) followed by in house created dose-response studies. This allowed us to determine potency (IC50s) and toxicity (CC50s) of each compound and focus on the 122 non-redundant compounds that had a selective index (SI) greater than 3 in Calu-3 cells (Fig 1a, Table S1). These compounds fall into a number of general categories with nucleosides accounting for 13% of the validated candidates.

**Figure 1.**
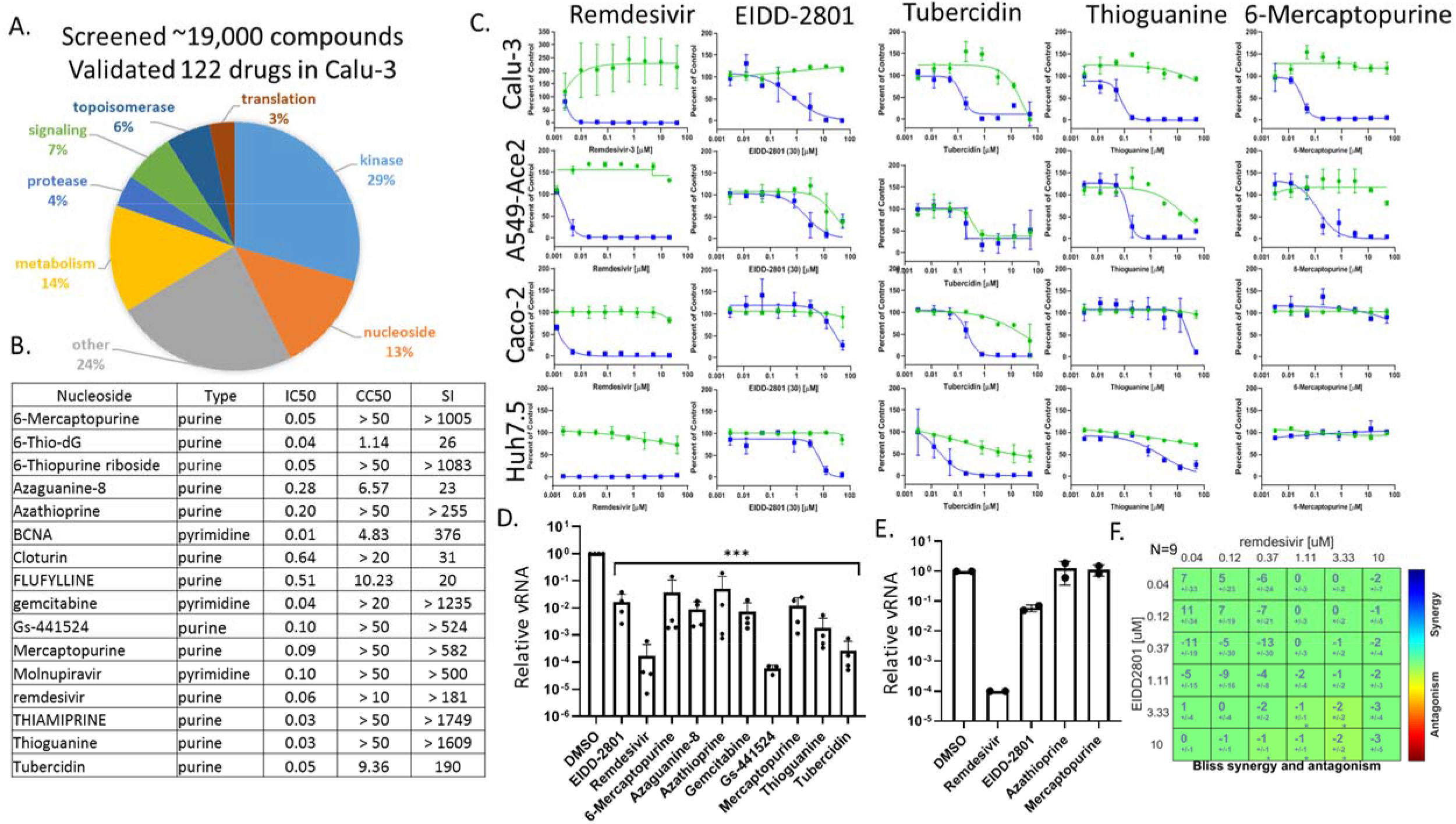
Antiviral nucleosides are highly active in respiratory cells and show cell-type specific activity. A. Pie chart of antivirals validated in Calu-3 cells with Selective Index (EC50/CC50)>3. From ∼19,000 compounds, 122 show activity. B. 16 nucleoside analogs validated in Calu-3 cells with the nucleoside type listed along with the EC50, CC50, and SI.C. Calu-3 A549-ACE2 Caco-2 Huh7.5 cells were treated with the indicated nucleosides in dose response showing infection (blue) and toxicity (green). D. Calu-3 cells pretreated with vehicle or 10uM of the indicated drugs and infected with SARS-CoV-2 for 48h and RT-qPCR analysis of viral infection with Mean±SE shown for reduction compared to vehicle control (n≥3). (p<0.001, ANOVA) E. RT-qPCR analysis of viral infection shown for the indicated drugs in nasal air-liquid interface cultures; Remdesivir (10uM), EIDD-2801 (10uM); Azathioprine (30uM); Mercaptopurine (30uM). (n=2).G. BLISS analysis of the 2×2 combination of Remdesivir and Molnupiravir in Calu-3 cells showing additivity.

Nucleoside analogs are a common class of drugs that are synthetic analogs that mimic their physiological counterparts and can be incorporated by cellular polymerases into DNA or RNA to inhibit cell division (*4, 13*). In addition, these can act as antimetabolites that deplete the supply of deoxynucleotides needed for DNA replication or nucleotides for RNA synthesis through inhibition of nucleoside biosynthesis enzymes. Nucleoside analogs and antimetabolites are generally used to treat cancer and for immunosuppression. A subgroup of nucleoside analogs are direct-acting antivirals as they are misincorporated by viral polymerases leading to defects in viral replication. Additional nucleoside analogs have been shown to have antiviral activity, likely through their antimetabolite activity. There are two characterized direct acting antiviral nucleosides shown to block SARS-CoV-2 infection in vitro and in vivo (*7, 14*). Remdesivir is FDA-approved for use in COVID-19, and is misincorporated by the viral RdRp leading to delayed chain termination (*15, 16*). Molnupirivir is in clinical trials, and is also misincorporated leading to increased viral mutagenesis and thus antiviral activity (*14, 17,18*). Importantly, we identified Remdesivir, the active metabolite of Remdesivir (Gs-441524), and Molnupiravir as antiviral in our screens (Fig 1a-c, Fig S1d). We identified 13 additional nucleoside analogs with antiviral properties against SARS-CoV-2 (Fig 1b, S2). To determine the breadth of antiviral activity of the nucleosides, we tested a panel of cell lines that are permissive to infection with SARS-CoV-2 including human respiratory A549-expressing human ACE2, human intestinal epithelial Caco-2 cells, human hepatocyte Huh7.5 and African green monkey Vero cells (Fig 1c, S1d). The known direct acting antivirals Remdesivir and Molnupiravir show activity across diverse cell types, with variable IC50s (Fig 1a-c, Fig S1d). We also found that the active form of Molnupiravir EIDD-1931 was active across cell types (Fig S1e). We validated antiviral activity of Remdesivir and Molnupiravir in Calu-3 cells using a quantitative RT-qPCR assay and found that these nucleoside analogs had antiviral activity by this assay (Fig 1d).

The additional antiviral nucleoside analogs we identified are thought to act as antimetabolites and are generally used for cancer or immunosuppression (*4*). We found that these drugs are non-toxic in Calu-3 cells at the antiviral concentrations, at least in part because these cells divide slowly. Moreover, we found that these nucleoside antimetabolites show very divergent cell type specific activity and toxicities (Fig 1a-d, Fig S1e). For example, tubercidin displays antiviral activity in Calu-3, Caco-2 and Huh7.5 cells with toxicity in A549-Ace2 and Vero cells. In contrast, Thioguanine and 6-Mercaptopurine are active in Calu-3 and A549-Ace2 cells but not active in Caco-2 or Vero cells. We also monitored the antiviral activity of a subset of these nucleoside analogs in Calu-3 cells by RT-qPCR, and found that they had significant antiviral activity (Fig 1d).We also tested a subset of these antiviral nucleosides in nasal air-liquid interface cultures and found that only Remdesivir and Molnupiravir showed potent antiviral activity as measured by RT-qPCR (Fig 1e). These studies demonstrate cell-type specific antiviral activities of nucleoside analogs that likely function as indirect inhibitors of nucleoside biology.

Given that the two direct acting antiviral nucleosides are distinct nucleoside derivatives, with Remdesivir an adenosine analog and Molnupiravir a cytosine analog, we tested whether the combination of these nucleosides would show antiviral synergy. Synergy is defined as having greater than an additive effect. Calu-3 cells were treated with six concentrations of each drug individually and in combination with each other resulting in a matrix of 36 drug-x drug-y concentration pairs monitoring viral infection. These assays were done as multiple biological and technical replicates, and we use BLISS analysis to determine if the two drugs interact(*19*)The null hypothesis in this model is that the drugs are additive, with a positive interaction leading to synergy and a negative interaction leading to antagonism. We found that the co-treatment of Calu-3 cells with Remdesivir and Molnupiravir during SARS-CoV-2 infection was additive (Fig 1f, Fig S3).

Antimetabolites are thought to act, at least in part, through inhibition of nucleoside biosynthesis (*4*). Cells have two pathways for nucleoside biogenesis, de novo synthesis and salvage pathways that recycle purines and pyrimidines from degradation products. While salvage pathways sustain cell viability, they cannot supply sufficient amounts of NTPs to allow for fast proliferation or support viral replication (*20*). Therefore, in addition to applications in cancer, the inhibition of the de novo nucleotide biosynthesis constitutes a broad-spectrum antiviral strategy. In support of this, our screen identified a subset of known inhibitors of host encoded nucleoside biosynthetic enzymes within the pyrimidine and purine biosynthetic pathways shown in Fig 2a. This included three pyrimidine biosynthesis inhibitors: the two DHODH inhibitors BAY-2402234 and Brequinar, as well as the UMPS inhibitor Pyrazofurin. Dose-response studies show that BAY-2402234 and Brequinar are active and have low toxicity in both Calu-3 and A549-ACE2 cells, and Pyrazofurin is active with low toxicity in Calu-3 cells (Fig 2b, 2c). We also discovered AVN944, a potent purine biosynthesis inhibitor active against IMPDH, is exhibiting antiviral activity at submicromolar concentrations with limited toxicity in Calu-3 cells, although AVN944 did not show selective activity in A549-ACE2 cells (Fig 2b, 2c). We did not identify the classical IMPDH inhibitor ribavirin in the screen that has been used to treat various RNA viruses, although ribavirin did inhibit viral infection at higher doses tested (>100uM) in Calu-3 cells (Fig. 2b, right most panel). We confirmed that the antiviral activity of the pyrimidine biosynthesis inhibitors was through nucleoside metabolism as we could fully block the antiviral activity of BAY-2402234, Brequinar or Pyrazofurin by treating Calu-3 cells with the pyrimidine nucleosides cytidine and/or uridine, but not the purine nucleosides adenosine or guanosine (Fig 2d). IMPDH is required for de novo guanosine biosynthesis and thus treatment of cells with the purine guanosine, but not adenosine or the pyrimidine nucleosides, could reverse the antiviral activity of AVN944 (Fig 2d). These data demonstrate that altering nucleoside pools can block viral replication.

**Figure 2.**
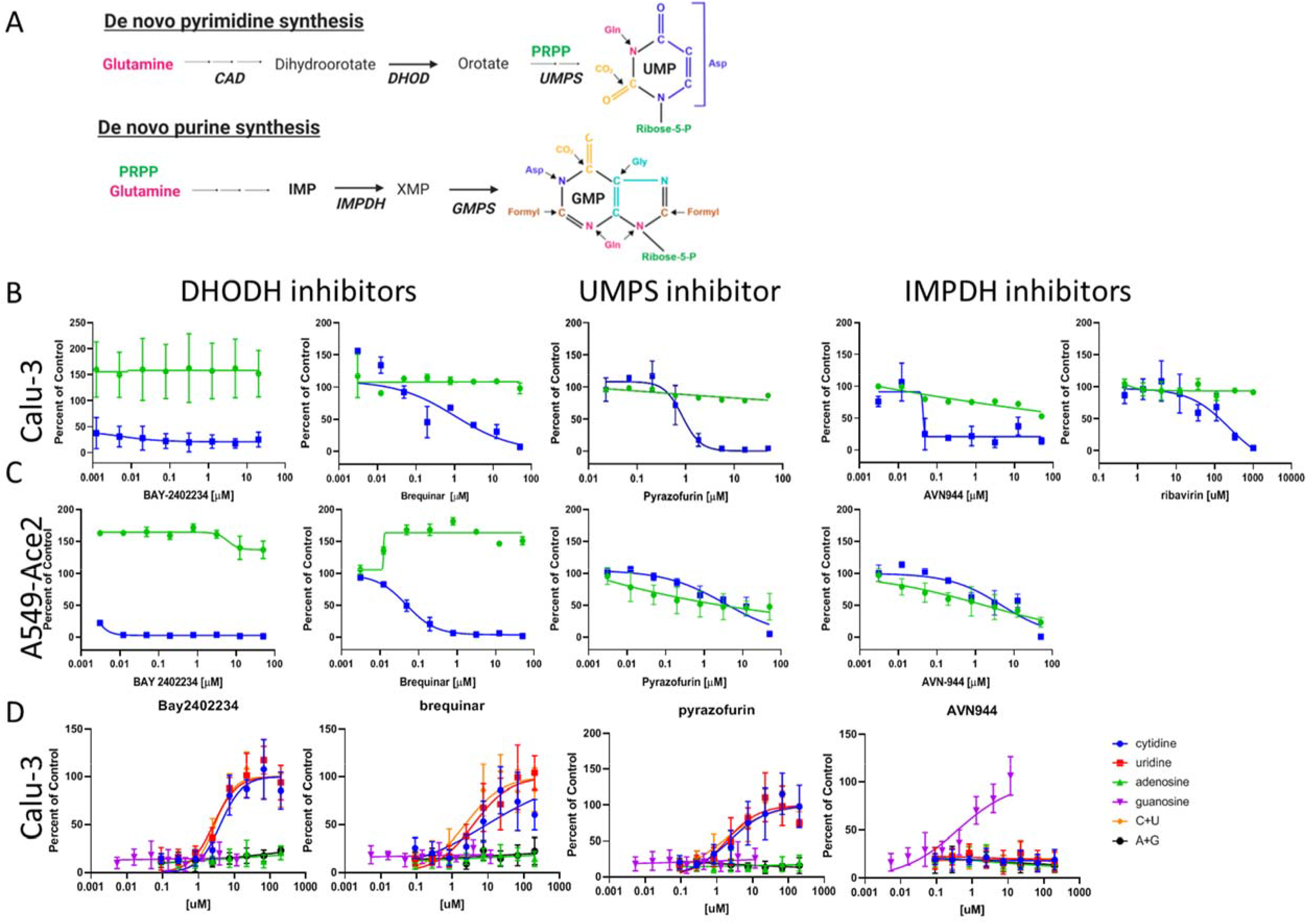
Inhibitors of host nucleoside metabolism are antiviral. A. Simplified schematic of nucleoside metabolism. (B-C). Dose response analysis of the pyrimidine biosynthesis or purine biosynthesis inhibitors in (B) Calu-3 cells or (C) A549-ACE2 cells. Infection (blue) and toxicity (green). (D) Analysis of Calu-3 cells treated with the indicated inhibitor in the presence of increasing concentrations of the indicated nucleosides.

Next, we set out to determine if altering pyrimidine or purine pools using host-directed metabolic inhibitors would synergize with the antiviral nucleoside analogs Remdesivir or Molnupiravir. Using our microscopy-based assay we performed 2×2 dose response analysis. We found striking synergy between DHODH inhibitors (Brequinar or BAY-2402234) and nucleoside analogs (Molnupiravir or Remdesivir) (Fig 3a-b, Fig S4-6). This synergy was observed in the sub-micromolar range of the DHODH inhibitor Brequinar and low nanomolar for BAY-2402234. In contrast, we observed little interaction between the IMPDH inhibitor AVN944 and either antiviral nucleoside analog, with a modest trend toward antagonism (Fig 3c). Given the emergence of SARS-CoV-2 variants and the promise of antivirals to have activity against diverse strains of SARS-CoV-2, we also tested whether these combinations would show antiviral synergy against a variant of SARS-CoV-2. We found that infection with the SARS-CoV-2 variant B.1.351 was inhibited by Brequinar, BAY-2402234, Pyrazofurin, AVN944, Molnupiravir and Remdesivir as single agents (Fig S7a). Moreover, we observed synergy between DHODH inhibitors (Brequinar or BAY-2402234) and nucleoside analogs (Molnupiravir or Remdesivir) but not the IMPDH inhibitor AVN944 (Fig S7b). This suggests that limiting the pyrimidine pool in combination with direct acting antivirals increases the antiviral activity of nucleoside analogs against diverse strains of SARS-CoV-2. In contrast, limiting guanosine has no increased benefit.

**Figure 3.**
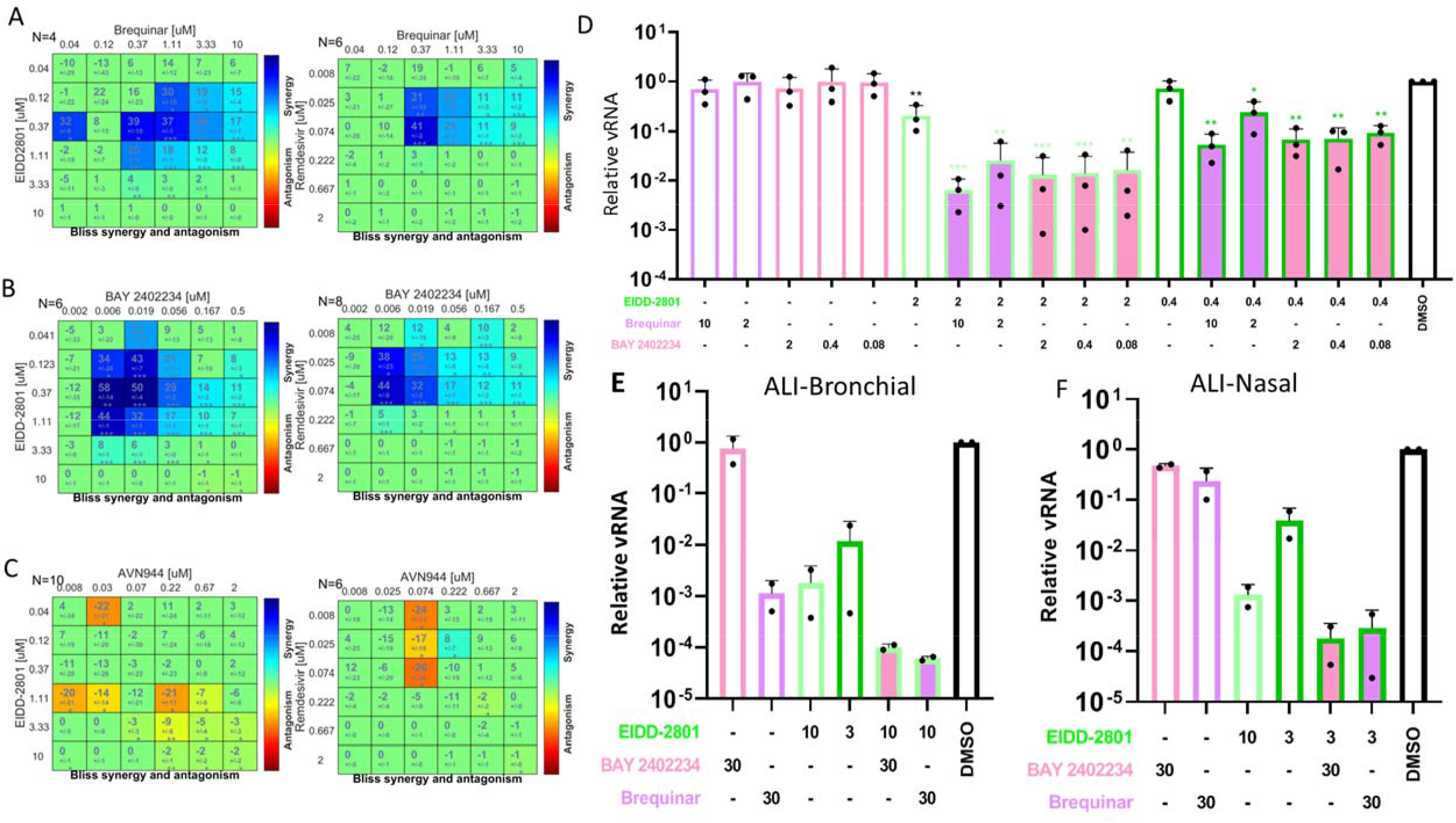
Combination of Molnupiravir or Remdesivir with DHODH inhibitors is synergistically antiviral in vitro. (A-C) BLISS analysis in Calu3 cells with Molnupiravir (EIDD-2801) or Remdesivir in combination with (A) DHODH inhibitor Brequinar; (B) DHODH inhibitor BAY-2402234; (C) IMPDH inhibitor AVN944. (D) Calu-3 cells were treated with the indicated drugs and infected with SARS-CoV-2. 48hpi viral replication was quantified by RT-qPCR and expression (viral RNA/18S) was normalized to vehicle treated cells. Mean±SE shown (n=3) Black asterisk show p-value relative to vehicle, ** p<0.01 ANOVA. Green asterisks show p-value relative to single treatment with Molnupiravir, * p<0.05, ** p<0.01, *** p<0.001, ANOVA. ALI-bronchial (E), or ALI-nasal (F) cells were treated with the indicated drugs and infected with SARS-CoV-2. 72hpi viral replication was quantified by RT-qPCR and expression (viral RNA/18S) was normalized to vehicle treated cells. Mean±SE shown (n=2). (D-F) Indicated concentration of drugs is in uM.

Therefore, we focused our downstream studies on DHODH inhibitors as they were active against SARS-CoV-2 in multiple cell types and Molnupiravir as it is an orally bioavailable drug and thus can be potentially used in an outpatient setting (*21, 22*). We confirmed that Molnupiravir shows dose-dependent inhibition in Calu-3 cells using a RT-qPCR assay (Fig 3d, Fig S8a). As single agents Brequinar and BAY-2402234 showed modest levels of inhibition as measured by RT-qPCR (Fig 3d). However, when we combined treatments, we observed striking decreases in viral replication upon co-administration of either of these DHODH inhibitors and Molnupirivir (Fig 3d).

We next tested the activity of these inhibitors in ALI respiratory epithelial cell cultures that more closely model the human respiratory epithelium. We used both tracheobronchial and nasal ALI cultures as these represent the two major sites of SARS-CoV-2 infection (*23*). We performed toxicity studies to identify doses of these drugs that did not impact epithelial barrier function (TEER), cilia beating frequency (CBF), or toxicity (LDH) in our nasal ALI cultures (*24*). We found that treatment with Molnuprivir, Brequinar, or BAY-2402234 at concentrations up to 30□ M were non-toxic in this system (Fig S8b). Therefore, we used these doses as the maximum in our ALI cultures. In bronchial ALI cultures, we found that Molnupiravir shows dose dependent activity and that Brequinar had significant single agent activity while BAY-2402234 had little activity as a single agent in these cells (Fig 3e). However, we found a significant reduction in viral replication upon co-treatment with either Brequinar or BAY-2402234 with Molnupiravir (Fig 3e). In the nasal cells, we found that Molnupiravir shows dose dependent activity and that neither Brequinar nor BAY-2402234 had significant single agent activity (Fig 3f). Again, we found a significant reduction in viral replication upon co-treatment of Molnupiravir with either DHODH inhibitor (Fig 3f).

Molnupiravir and Brequinar are both orally dosed drugs that are undergoing clinical trials in COVID-19 patients, and we observed synergistic antiviral activity with these drugs in diverse model cell systems. Therefore, we set out to test whether the combination of Molnupiravir and Brequinar would show benefit in the treatment of SARS-CoV-2 infection in vivo. We used a mouse model of infection where wild type Balb/C mice are intranasally inoculated with SARS-CoV-2 strain B.1.351 (1e5 PFU/mouse). In this model we found robust replication 2dpi as measured by viral titers in the lungs (∼10^8 PFU/g lung), bronchiolar sloughing of infected epithelial cells, and significant inflammatory cell infiltration including edema with peribronchiolar and perivascular cuffing as measured by histology (Fig 4). As expected, when we treated the animals with Molnupiravir (EIDD-2801) we observed dose-dependent reduction in viral titers (Fig 4a). In addition, there was only minor mitigation of the lung pathology with significant epithelial cell infection and inflammatory cell infiltration other than at the highest dose of Molnupiravir tested (150mg/mL) (Fig 4b). We also treated mice with Brequinar alone and observed no impact on viral titers or lung pathology at 2 days post infection (Fig 4a-b). Next, we combined treatments with multiple dosing of Molnupiravir and a single dose of Brequinar (20mg/kg). We found a ∼4 Log reduction in viral titers upon co-treatment with Brequinar and Molnupiravir at the highest dose of Molnupiravir, and a significant reduction in titers at both 150 mg/kg and 50 mg/kg Molnupiravir when combined with Brequinar compared to Molnupiravir alone (Fig 4a). Strikingly, we observed strong suppression of inflammation in the lung where even at the lowest combination dose there is reduced peribronchiolar and perivascular cuffing with lessened alveolar and interstitial inflammation and edema (Fig 4b). Moreover, co-treatment led to a clear protection of lung architecture with little alterations to bronchiolar and alveolar cells. Altogether, this combination of Molnupiravir and Brequinar for COVID-19 treatment shows promise as we observe both reduction in viral replication and decreased pathology. We also performed an experiment where we used a single dose of Molnupiravir (50mg/kg) alone, and in combination with increasing doses of Brequinar (20mg/kg and 50mg/kg) (Fig 4c-d). We again observed significant decreases in viral titers upon co-treatment of Molnupiravir and Brequinar over Molnupiravir alone (Fig 4c). Histology analysis revealed that treatment with Molnupiravir modestly reduced SARS-CoV-2-induced pathology and that co-administration with Brequinar further reduced the inflammatory response seen with Molnupiravir alone (Fig 4d).

**Figure 4.**
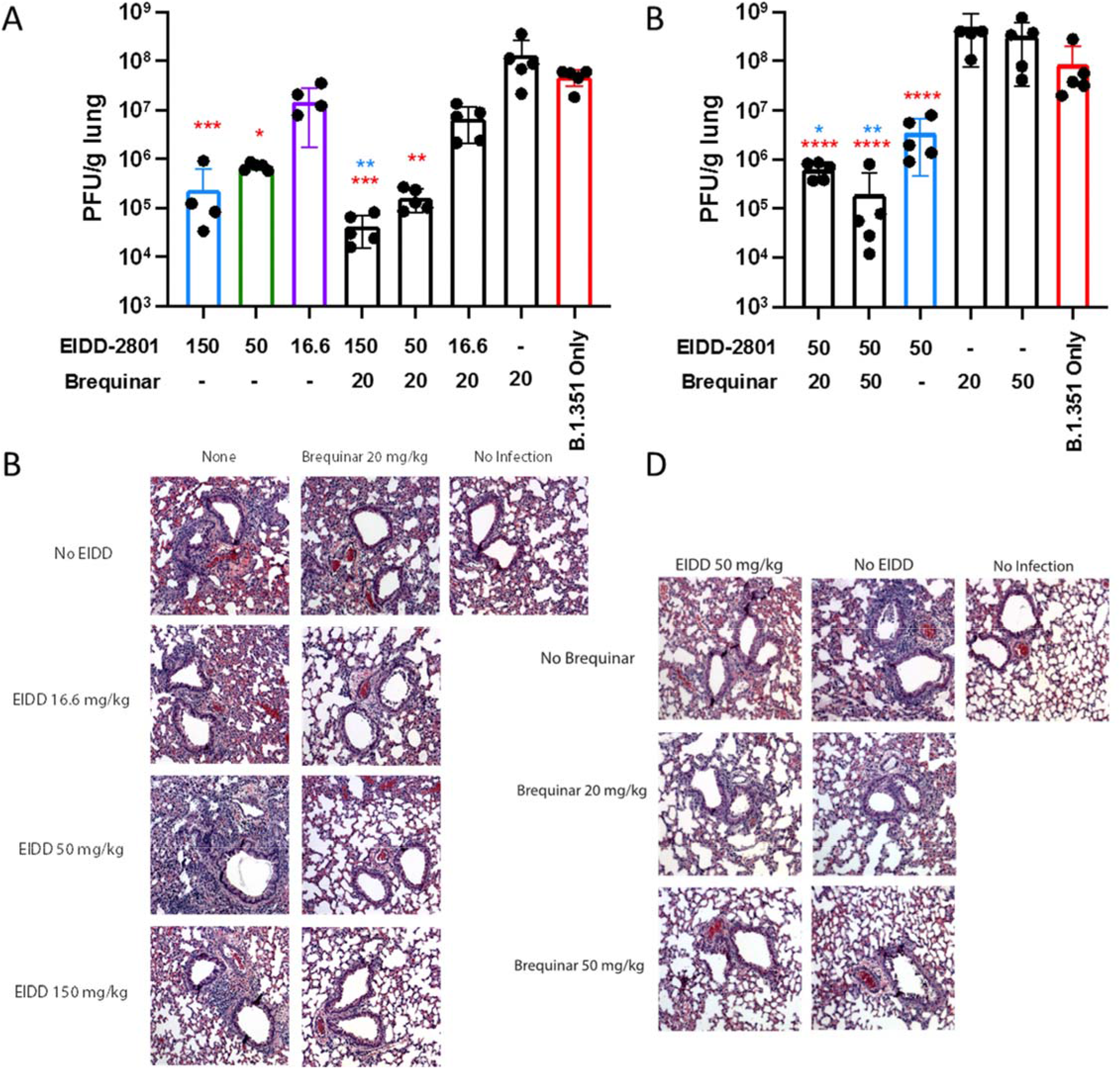
Combination of Molnupiravir and Brequinar reduces SARS-CoV-2 infection and inflammation in vivo. Wild type Balb/C mice were treated with the Brequinar (IP) and/or EIDD-2801 (PO) daily at the indicated concentrations (mg/kg) starting at 12 hours before infection. Mice were intranasally inoculated with 1×10^5^ PFU/mouse of SARS-CoV-2/B.1.351. At 2 dpi, mice were euthanized and lungs analyzed for viral titer by (A,C) plaque assay or (B,D) fixed in 4% PFA for H&E staining. N=5 mice per group. Plaque assay titer calculations are reported as One-way Anova with Dunnett multiple comparisons test. *=p<0.05.

Given that for many viral infections combinations of antivirals are needed to suppress infection, we suggest that combining nucleoside analogs with DHODH inhibitors would be beneficial, as the combination would both reduce replication and inflammation due to SARS-CoV-2 infection within therapeutic ranges.

## Acknowledgments

We thank members of the Cherry, Lynch, Frieman and Penn Center for Precision Medicine for advice and discussion. We thank all members of the High-Throughput Screening Core at University of Pennsylvania for reagents and technical support. We thank S. Weiss and Y. Li for sharing SARS-Related Coronavirus 2, Isolate USA-WA1/2020 and Isolate B.1.351 (obtained from the Centers for Disease Control and BEI resources) and Andy Pekosz for B.1.351 for mouse studies. We thank Xin Hu, Richard Eastman, Matthew Hall, and members of NCATS Compound Management for assembly and preparation of the NCATS compound collections. We thank CALIBR for providing the ReFrame library and validation plates.

## Author contributions

Research design D.C.S, R.M.J., E.L. S.C. M.F, M.B.F. and S.C.; experimentation D.C.S, R.M.J., K.A., J.M, K.W.,B.K.,M.D.,S.W.,H.L.H.,C.D.,L.C.,J.S.L.,M.L., E.L. S.C. M.F, M.B.F. and S.C; data analysis D.C.S, R.M.J., K.A., J.M, K.W., B.K., M.D., S.W., H.L.H., C.D., L.C., J.S.L., M.L., E.L., S.C. M.F, M.B.F. and S.C; writing D.C.S., E.L., S.C. M.F, C.A.T., M.B.F. and S.C; supervision D.C.S. S.C. M.F., M.B.F. and S.C.

## Data and materials availability

All data in this paper are presented in the main text and supplementary text.

