## Supplemental Data for "Pyrimidine biosynthesis inhibitors synergize with nucleoside analogs to block SARS-CoV-2 infection"

**Supplementary Materials:**

**<u>Methods</u>**:

<u>Viruses and cells:</u>

For cell culture studies, SARS-CoV-2 was obtained from BEI (USA/WA-1 strain, B.1.351 strain). SARS-CoV-2 was amplified in Vero CCL81 to create a P1 stock at $1.5x10^6$ TCID50/mL, was sequence verified, and used for all cell culture experiments. All work with infectious virus was performed in a Biosafety Level 3 laboratory and approved by the Institutional Biosafety Committee and Environmental Health and Safety.

Vero (ATCC, CCL81) and Huh7.5 cells (C. Rice, Rockefeller) were cultured in DMEM, supplemented with 10% (v/v) fetal bovine serum, 1% (v/v) penicillin/streptomycin, 1% (v/v) L-Glutamax. Calu-3 cells (ATCC, HTB-55) were cultured in MEM, supplemented with 10% (v/v) fetal bovine serum, 1% (v/v) penicillin/streptomycin, 1% (v/v) L-glutamine, and 1% (v/v) non-essential amino acids. A549-ACE2 cells were cultured in RPMI1640 supplemented with 10% (v/v) fetal bovine serum, 1% (v/v) penicillin/streptomycin, 1% (v/v) L-Glutamax. Caco-2 (ATCC, HTB-37) were cultured in MEM alpha supplemented with 20% (v/v) fetal bovine serum, 1% (v/v) penicillin/streptomycin, 1% (v/v) L-glutamine. All cells were grown at 37°C, 5% $CO_2$ and 20% $O_2$.

Day 20 air liquid interface EpiAirway tracheobronchial tissues were obtained from MatTek Corporation (AIR-100) and used after Day 30. Tissues were fed twice a week until use. Air liquid interface pooled nasal epithelial cultures from Epithelix were fed twice a week until use. Toxicity studies were performed on nasal air-liquid interface cultures. Daily basolateral exposure of test compounds was evaluated using tissue integrity (TEER); cytotoxicity as basolateral LDH release and cilia beating frequency (CBF) at 72h.

*Tissue Integrity (TEER)* Tissue integrity is determined by monitoring trans-epithelial electrical resistance (TEER) using the EVOM3 Ohm Meter (World Precision Instruments). Resistance values (Ω) were converted to TEER ($\Omega.cm^2$) using the following formula: TEER ($\Omega.cm^2$) = (resistance value (Ω) - 100(Ω)) x 0.33 ($cm^2$), where 100 Ω is the resistance of the membrane and 0.33 $cm^2$ is the total surface of the epithelium.

*Cytotoxicity (LDH release)* Basolateral lactate dehydrogenase release was quantified using Cytotoxicity LDH Assay Kit-WST (Dojindo, CK12-20), measuring the absorbance of each sample at 490nm with a microplate reader. To determine the percentage of cytotoxicity, the following equation was used (A = absorbance values): Cytotoxicity (%) = (A (exp value)-A (low control)/A (high control)-A (low control))*100. The high control value was obtained by 10 % Triton X-100 apical treatment (24 hours). Triton X-100 causes a massive LDH release and corresponds to 100 % cytotoxicity.

*Cilia Beating Frequency (CBF)* Cilia beating frequency was measured using the following setup: a Sony XCD V60 camera connected to an Olympus BX51 microscope with a 5x objective and a camera specific software. The cilia beating frequency is expressed as Hz. 256 images were captured at high frequency rate (125 frames per second) at 34°C, cilia beating frequency was then calculated using Cilia-X software (Epithelix).

For cell culture studies, SARS-CoV-2 was obtained from BEI (USA/WA-1 strain, B.1.351 strain). SARS-CoV-2 was amplified in Vero CCL81 to create a P1 stock at $1.5x10^6$ TCID50/mL, was sequence verified, and used for all cell culture experiments. All work with infectious virus was performed in a Biosafety Level 3 laboratory and approved by the Institutional Biosafety Committee and Environmental Health and Safety.

High-Throughput Screening:

Ten-thousand Calu-3 cells were plated per well of 384 well assay plates (Corning ) in 20ul of growth medium. For the PENN and NCATS libraries, 50nL of drugs were added at a final concentration of 0.8 µM in 0.2% DMSO. For the ReFrame library, 20ul of MEM was dispensed to compound source plates pre-spotted with 50nL of 10mM compound in DMSO.  5ul of diluted compound was added per well of 384W assay plate yielding a final concentration of 3.6uM in 0.04% DMSO. The positive control 10uM Remdesivir (n=32) and the negative control 0.2% DMSO (n=32) were spotted on each plate. One hour post drug addition, cells were infected with SARS-CoV-2 (MOI=0.5). Cells were fixed 48hpi in 4% formaldehyde/PBS for 15min at room temperature, washed three times with PBS, blocked with 2% BSA/PBSTfor 60 minutes, and incubated in primary antibody (anti-dsRNA J2) overnight at $4^oC$. Cells were washed 3x in PBST with an automated plate washer (Biotek) and incubated in secondary antibody (anti-mouse alexa 488 and Hoescht 33342) for 1h at room temperature. Cells were washed 3x in PBST with an automated plate washer and imaged using an automated microscope (ImageXpress Micro, Molecular Devices). Cells were imaged with a 10X objective, and four sites per well were captured. The total number of cells and the number of infected (dsRNA+) cells were measured using the cell scoring module (MetaXpress 5.3.3), and the percentage of infected cells was calculated.  The aggregated percent infection of the 0.2% DMSO (n=32) and 10uM Remdesivir control wells (n=32) on each assay plate were used to calculate z'-factors, as a measure of assay performance and data quality. Sample well infection was normalized to aggregated DMSO plate control wells and expressed as Percentage of Control [POC = ($\%Infection_{sample}$/ Average $\%Infection_{DMSO}$)*100] and Z-score [Z= ($\%Infection_{sample}$ - Average $\%Infection_{DMSO}$) / Standard Deviation $\%Infection_{DMSO}$]in Spotfire (PerkinElmer). Candidate hits were selected as the following: for UPENN screen compounds with POC <40% for infection and viability >80% in either replicate; for NCATS screen POC <40% for infection and viability >80%; For the ReFrame screen, POC <50% for infection and viability >60% viability for the average of the replicates, compared to DMSO control.

Dose response studies:

Candidate drugs from the UPENN and NCATS library were purchased as powders from Selleckchem, MedchemExpress, Cayman and MedKoo and suspended in DMSO. Drugs were arrayed in 8-pt dose-response in 384 well plates. For the ReFrame validation, 15ul of MEM was dispensed to compound source plates pre-spotted with 100nL of compound in DMSO arrayed in an 8 pt dose response. 5ul of diluted compound was added per well of 384W assay plate yielding final concentrations of 9.5, 3.1, 1.0, 0.35, 0.12, 0.04, 0.01, and 0.004 uM in 0.1% DMSO. Calu-3 (10,000 cells), A549-ACE2 (3,000), Caco-2 (1,500), Huh7.5 (3,000) or Vero (3,000) were plated in 384 well plates. Twenty-four hours post plating (72 hours for Caco-2), drug additions and infections were performed using the screening conditions. 0.2% DMSO (n=32) and 10 µM Remdesivir (n=32) were included on each plate as controls for normalization.  For nucleoside rescue experiments, cells were treated with an EC90 concentration of a given nucleoside biosynthesis inhibitor in combination with a dose response of nucleosides. Infection at each drug concentration was normalized to aggregated DMSO plate control wells and expressed as percentage-of-control (POC=% Infection $_{sample}$/Avg % Infection $_{DMSO\ cont}$). A non-linear regression curve fit analysis (GraphPad Prism 8) was performed on POC of % Infection and cell viability using log10 transformed concentration values to calculate IC50 values for % Infection and CC50 values for cell viability for each drug/cell line combination. The IC50 and CC50 values represent the average of ≥ 2 independent experimental replicates. Selectivity index (SI) was calculated as a ratio of drug's CC50 and IC50 values (SI = CC50/IC50).

Quantitative drug combination analysis:

Using our HTS assay conditions, drugs were added to Calu-3 cells in 384W assay plates using a Tecan D300e dispenser in a matrix that combined six concentrations of Remdesivir (up to 2µM) or Molnupiravir (up to 10uM) with six concentrations of Brequinar (up to 10uM), BAY-2402234 (up to 0.5uM), and AVN944 (up to 2uM), resulting in 36 concentration combinations per drug pair in 0.2% DMSO. Each combination was independently repeated at least 3 times as technical duplicates in each biological replicate. Sample well infection was normalized to aggregated 0.2% DMSO plate control wells (n=32) and expressed as Percentage of Control. Synergy between drug combinations was determined by the Bliss independence model, to quantitatively assess drug interaction patterns within the drug-drug combination matrix (*1, 2*) .The Bliss expectation (E) for a combined response was calculated by $E = (A + B) - (A \times B)$ where A and B are the fractional inhibition of SARs-CoV-2 infection of drug A and B at a given dose. The difference between the Bliss expectation and the observed inhibition of SARS-CoV-2 infection for the combination of drugs A and B at the same dose is the BLISS value. Bliss values between 0-10 indicate that the combination is additive (as expected for independent pathway effects); Bliss values >20 indicates activity greater than additive (synergy); and values <0 indicate the combination is less than additive (antagonism).

RT-qPCR:

Calu-3 cells (750,000 cells/well) were plated in 6 well plates. The next day, drugs were added to cells. One hour later cells were infected with SARS-CoV-2 (MOI=0.3). For ALI cultures, the apical surface was washed with OptiMEM, and cells placed into fresh media with drugs added to the basolateral surface. Cells were infected apically with SARS-CoV-2 (MOI=0.2) for one hour and subsequently the virus inoculum was removed. The cells were placed into fresh media daily with the indicated drugs. At the indicated time point, total RNA was purified using Trizol (Invitrogen) followed by RNA Clean and Concentrate kit (Zymo Research) 48 hpi for Calu-3 and ALI cultures. For cDNA synthesis, reverse transcription was performed with random hexamers and Moloney murine leukemia virus (M-MLV) reverse transcriptase (Invitrogen). Synthesized RNA was used as a standard (BEI). Gene specific primers to SARS-CoV-2 (Wuhan v1, NSP14) and SYBR green master mix (Applied Biosystems) were used to amplify viral RNA and 18S rRNA primers were used to amplify cellular RNA using the QuantStudio 6 Flex RT-PCR system (Applied Biosystems). Relative quantities of viral and cellular RNA were calculated using the standard curve method (*3*). Viral RNA was normalized to 18S RNA for each sample (Wuhan V1/18S) (*4*).

Mouse studies:

SARS-CoV-2 B.1.351 was generously provided by Dr. Andy Pekosz at The Johns Hopkins University School of Public Health. Stock virus was prepared by infection of Vero E6/TMPRSS2 cells in growth media (DMEM (Quality Biological), supplemented with 10% (v/v) fetal bovine serum (Gibco), 1% (v/v) penicillin/streptomycin (Gemini Bio-products) and 1% (v/v) L-glutamine (2 mM final concentration, Gibco) fetal bovine serum plus for two days when CPE was visible. Media were collected and clarified by centrifugation prior to being aliquoted for storage at −80°C. Titer of stock was determined by plaque assay using Vero E6 cells as described previously (*5*). All work with infectious virus was performed in a Biosafety Level 3 laboratory and approved by The University of Maryland School of Medicine Institutional Biosafety Committee. Mouse challenge studies were approved by The University of Maryland School of Medicine IACUC. EIDD-2801 (MedChemExpress #HY-135853) was resuspended in corn oil (Sigma #8267) and 10% DMSO (Sigma #2438), with dosing twice a day as oral gavage at 150mg/kg. Brequinar (MedChemExpress #HY-108325) was resuspended in 10% DMSO and sterile saline, with dosing once a day as intraperitoneal injection.

Mice were anaesthetized by intraperitoneal injection with 50 μL of a mix of xylazine (0.38 mg/mouse) and ketamine (1.3 mg/mouse) diluted in phosphate buffered saline (PBS). Mice were intranasally inoculated with $1 \times 10^5$ pfu of B.1.351 variant of SARS-CoV-2 in 50 μL. Challenged mice were weighed on day of infection and daily for 2 days post infection. 2-days post infection, 5 mice were sacrificed from each treatment and control group, lungs were harvested to determine viral titer by a plaque assay, and fixed in 4% paraformaldehyde for 24 hours before sectioning and staining with hematoxylin and eosin by UMSOM Histology Core(*5*).

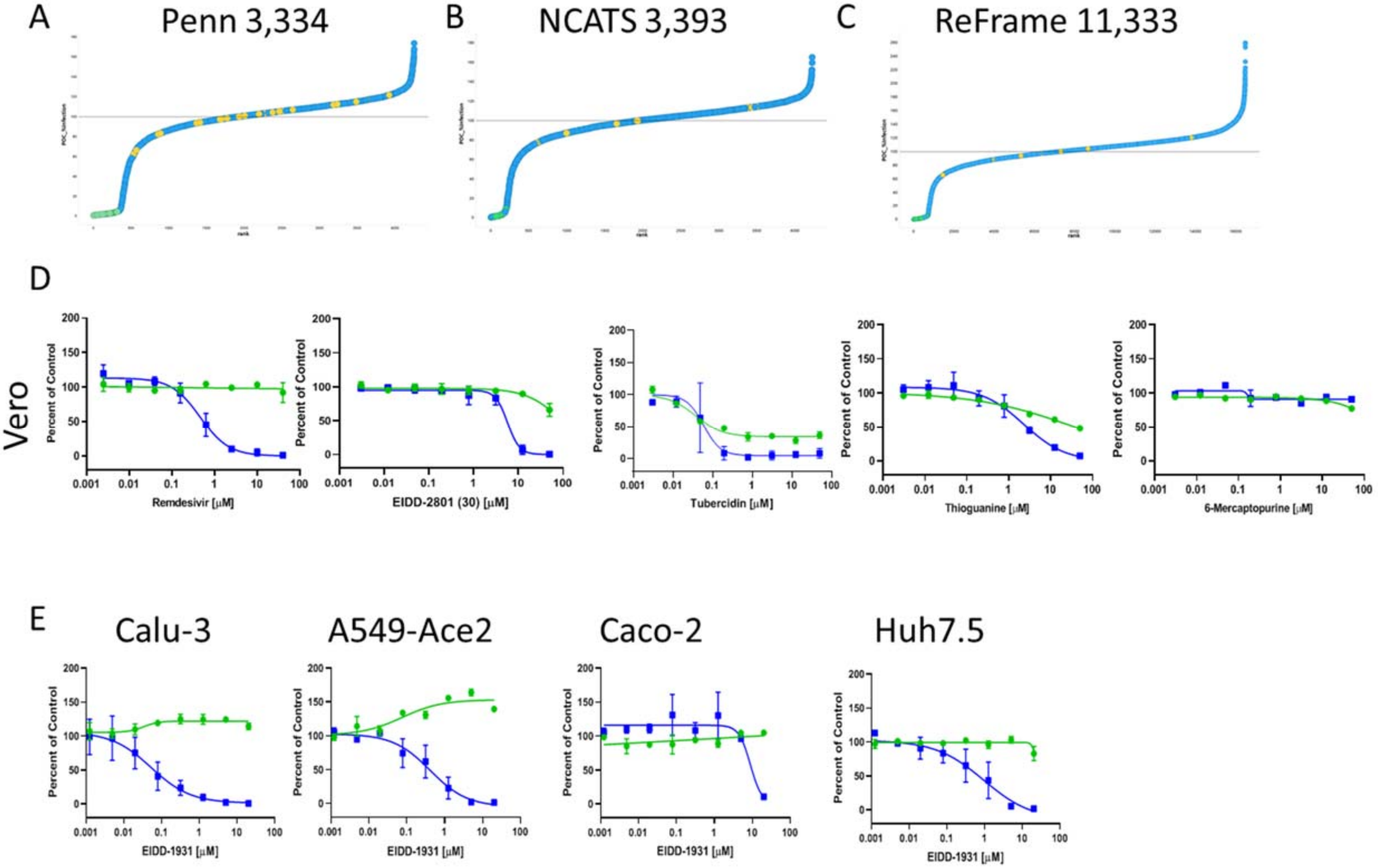


**Figure S1. High-throughput screening in Calu-3 cells to identify antivirals against SARS-CoV-2.**

A. High throughput screening of ~18,000 drugs including the UPENN library, NCATS library and ReFrame library in Calu-3 cells infected with SARS-CoV-2. Percent of Control (POC) for % infection is plotted versus rank of the drug across the primary screens. Yellow circles, DMSO controls; Green circles, remdesivir controls; Blue Circles, sample wells. Solid horizontal line represents 100% Infection of DMSO control wells. D. Vero cells were treated with the indicated nucleosides in dose response showing infection (blue) and toxicity (green). E. The indicated cell line was treated with increasing doses of EIDD-1931 and dose responses for infection (blue) and toxicity (green) are shown.

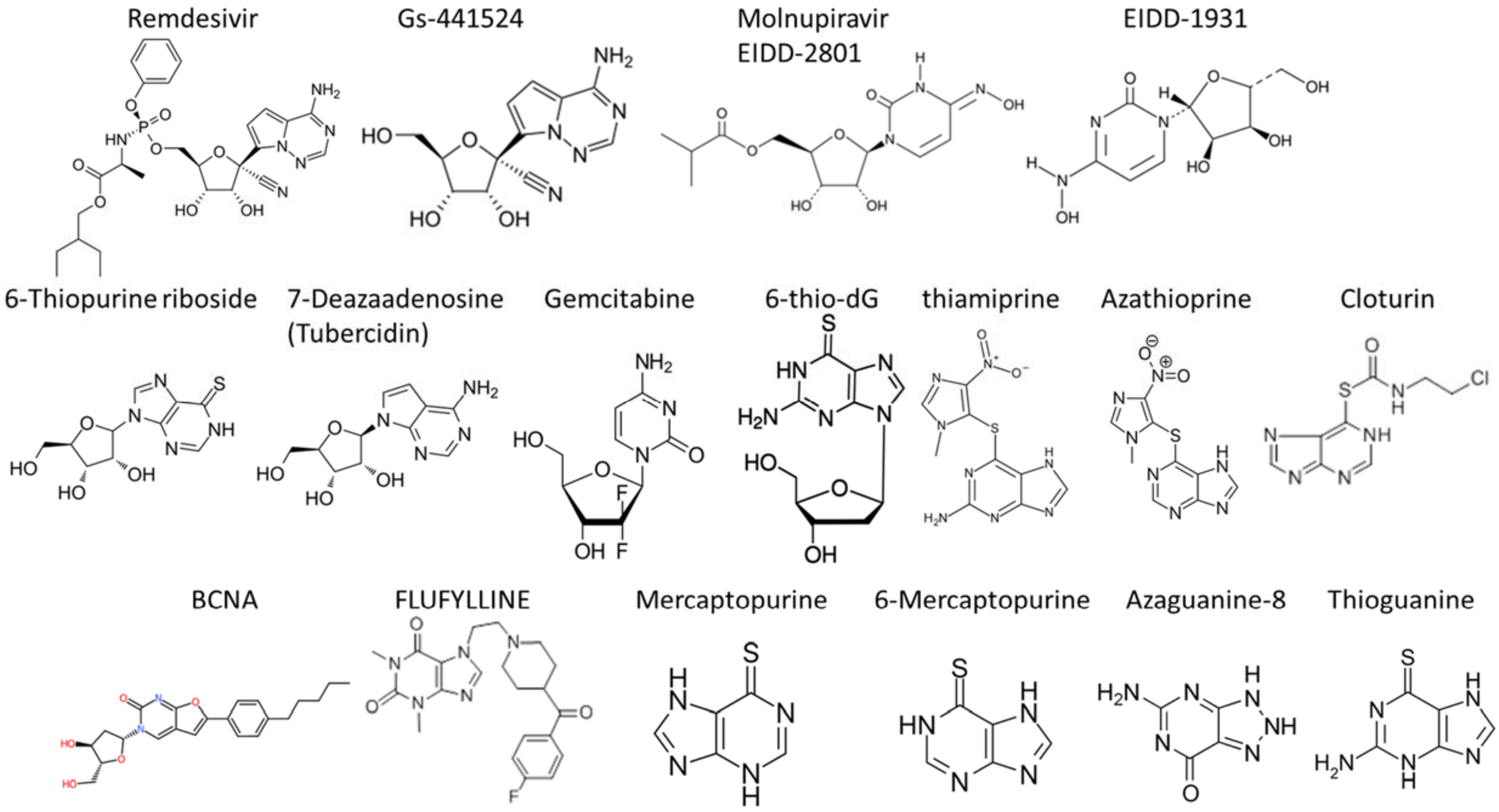


**Figure S2. Structures of antiviral nucleoside analogs shown.**

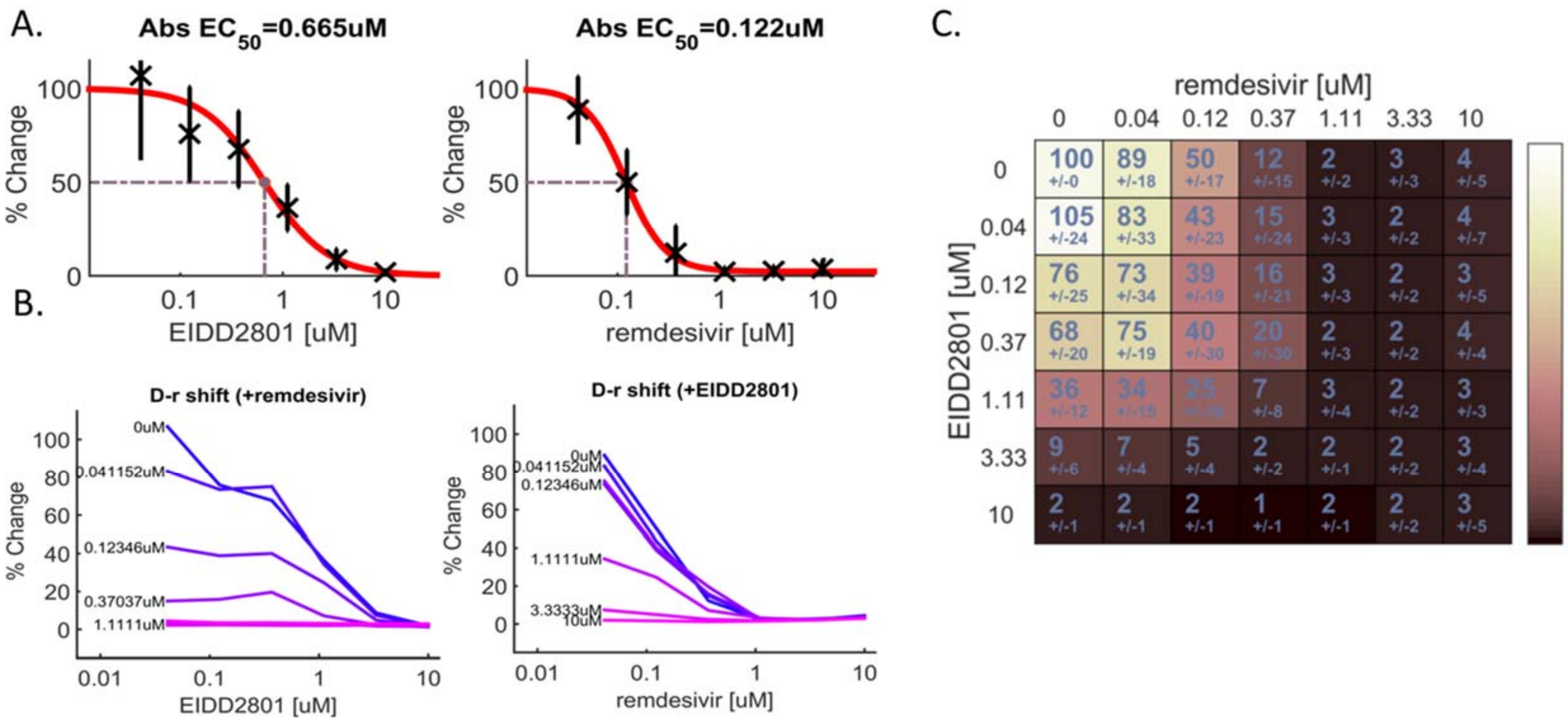


**Figure S3. Combination of Remdesivir and Molnupiravir is additive.** A. anti-SARS2 activity of Remdesivir and Molnupiravir as single agents. B. anti-SARS2 activity of Remdesivir in combination with a single concentration of Molnupiravir or Molnupiravir in combination with a single concentration of Remdesivir. C. Two-dimensional representation of dose response interaction matrix for percent of control of infection.

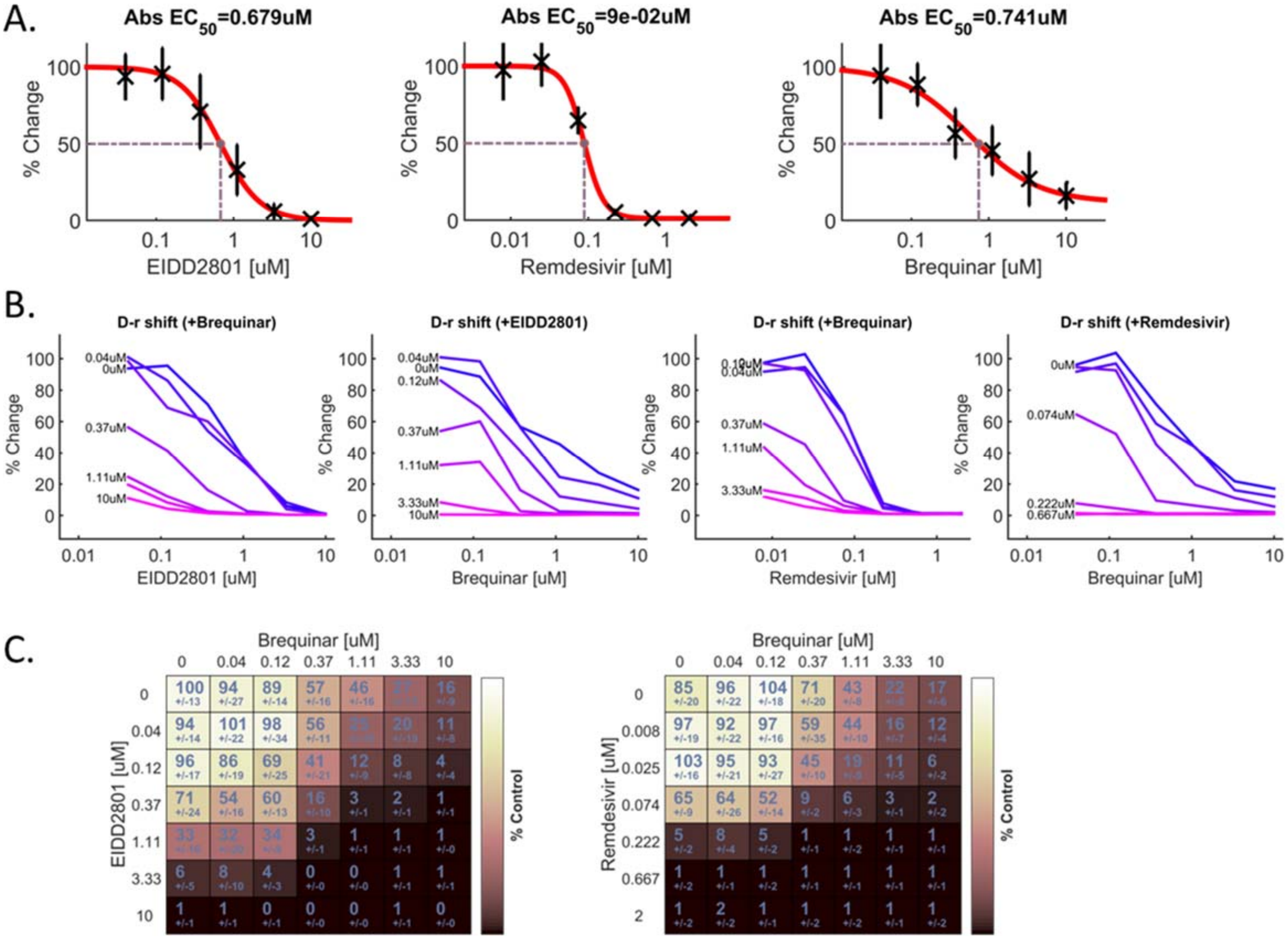


**Figure S4 Combination of Molnupiravir or Remdesivir with DHODH inhibitor Brequinar is synergistically antiviral in vitro.** A. anti-SARS2 activity of Molnupiravir, Remdesivir, and Brequinar, as single agents. B. anti-SARS2 activity of molnupiravir or remdesivir in combination with a single concentration of Brequinar. C. Two-dimensional representation of dose response interaction matrix for percent of control of infection.

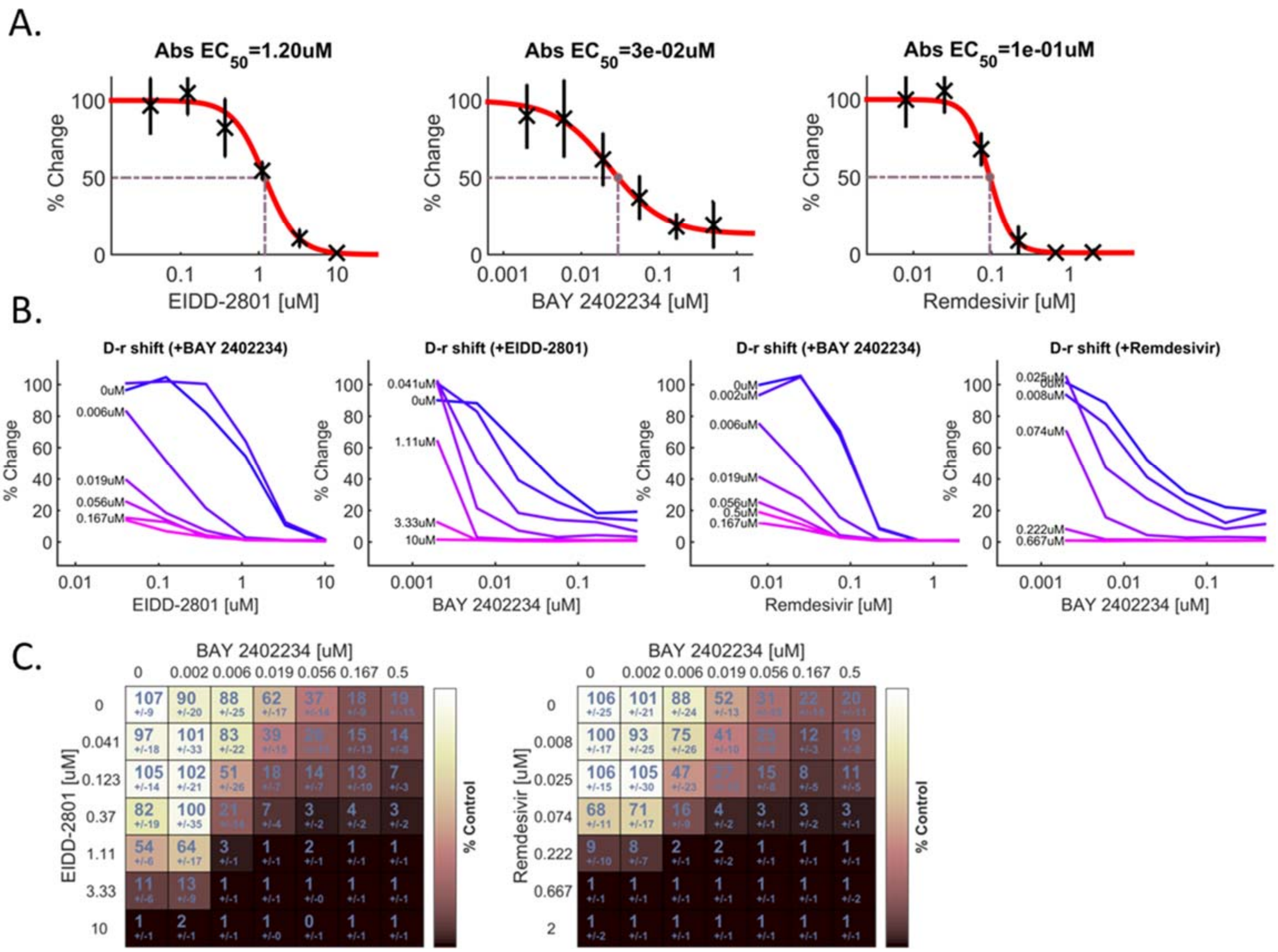


**Figure S5. Combination of Molnupiravir or Remdesivir with DHODH inhibitor BAY-2402234 is synergistically antiviral in vitro.** A. anti-SARS2 activity of Molnupiravir, Remdesivir, and BAY-2402234 as single agents. B. anti-SARS2 activity of Molnupiravir or Remdesivir in combination with a single concentration of BAY-2402234. C. Two-dimensional representation of dose response interaction matrix for percent of control of infection.

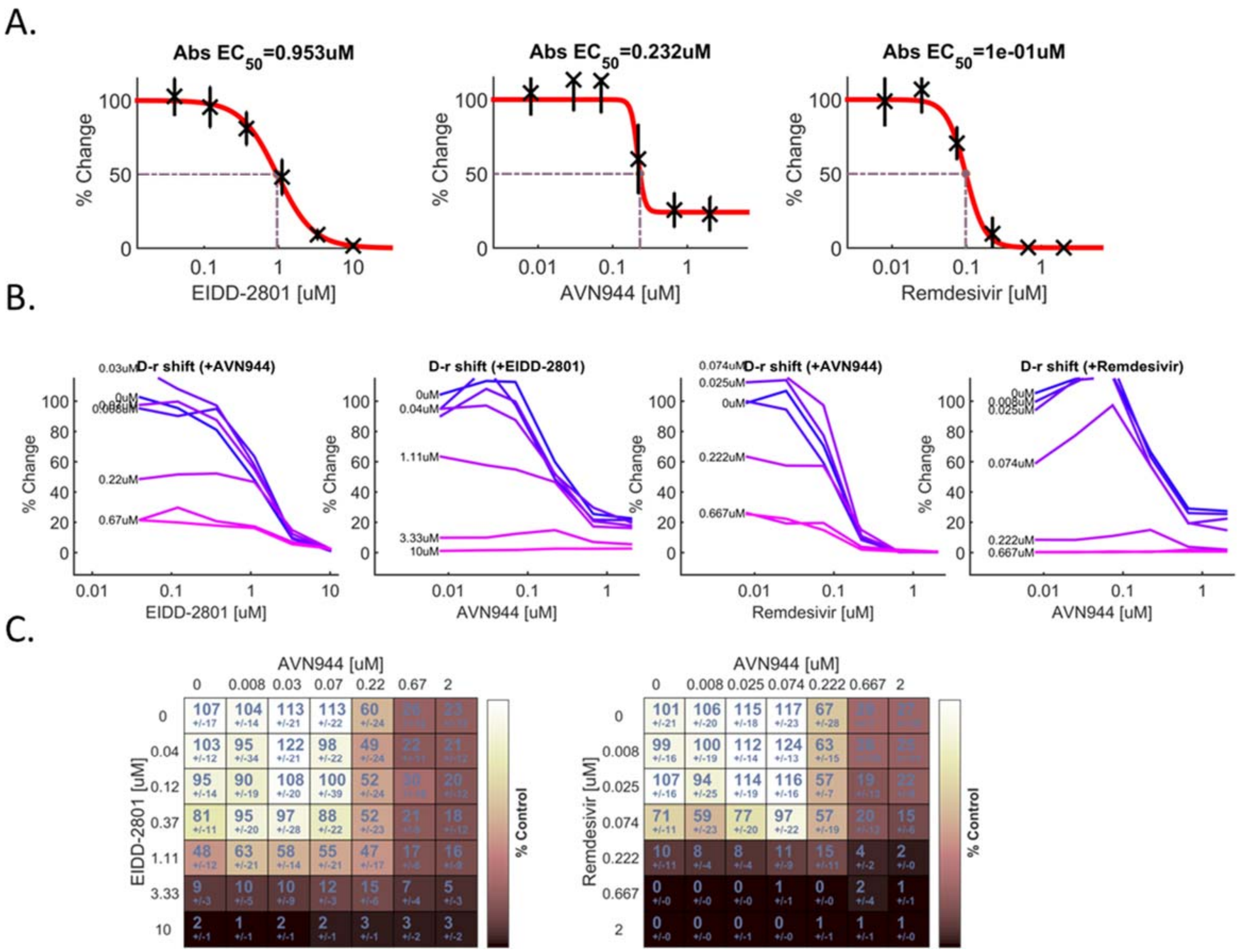


**Figure S6. Combination of Molnupiravir or Remdesivir with IMPDH inhibitor AVN944 in vitro.** A. anti-SARS2 activity of Molnupiravir, Remdesivir, and AVN944 as single agents. B. anti-SARS2 activity of Molnupiravir or Remdesivir in combination with a single concentration of AVN944. C. Two-dimensional representation of dose response interaction matrix for percent of control of infection.

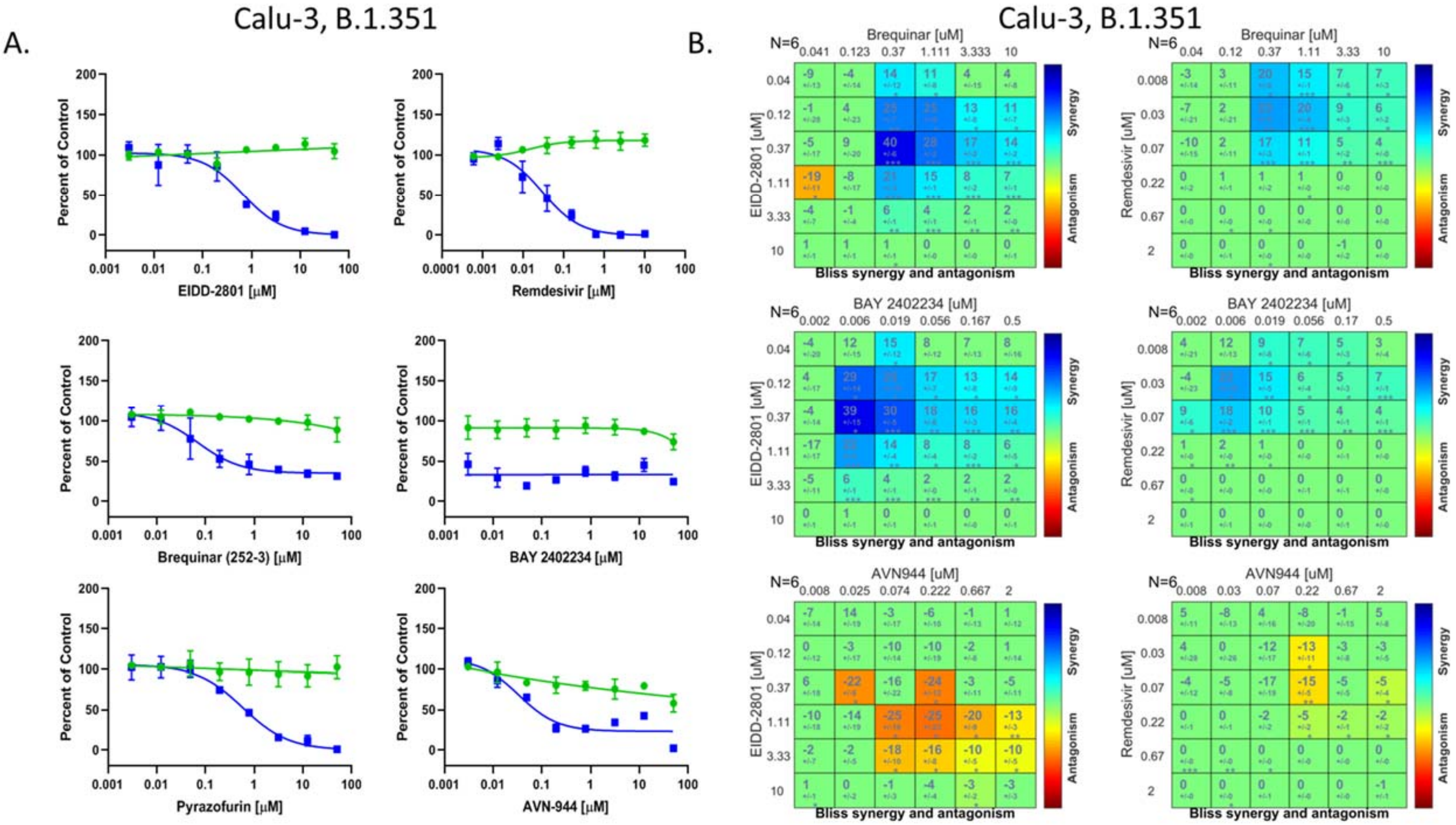


**Figure S7. Combination of Molnupiravir or Remdesivir with DHODH but not IMPDH inhibitor is synergistic against SARS-CoV-2 B.1.351 in vitro.** A. Dose response analysis of the pyrimidine biosynthesis or purine biosynthesis inhibitors in Calu-3 cells infected with SARS-CoV-2 B.1.351. Infection (blue) and toxicity (green). B. BLISS analysis in Calu3 cells with Molnupiravir (EIDD-2801) or Remdesivir in combination with DHODH inhibitor Brequinar; DHODH inhibitor BAY-2402234 or IMPDH inhibitor AVN944 infected with SARS-CoV-2 B.1.351.

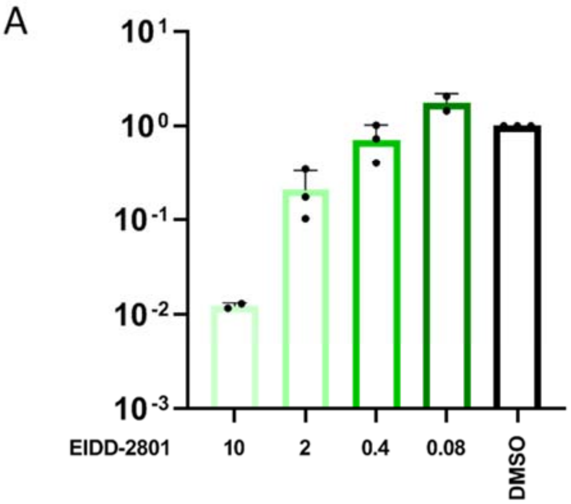


B

Maximal tested tolerated dose in nasal air-liquid interface cultures at 72h post basolateral daily exposure

| Compound | TEER | CBF | LDH |
| --- | --- | --- | --- |
| Molnupiravir | 30 uM | 30 uM | 30 uM |
| Remdesivir | 30 uM | 30 uM | 30 uM |
| Brequinar | 30 uM | 30 uM | 30 uM |
| BAY2402234 | 30 uM | 30 uM | 30 uM |
| AVN944 | 30 uM | >10 uM | 10 uM |

**Figure S8. Nucleoside-related drugs are well-tolerated in air-liquid interface cultures.** A. Calu-3 cells were treated with the indicated concentrations of Molnupiravir (EIDD-2801) and infected with SARS-CoV-2. 48hpi viral replication was quantified by RT-qPCR and expression (viral RNA/18S) was normalized to vehicle treated cells. Mean±SE shown. B. Nasal air-liquid interface cells were treated daily on the basolateral side with the indicated concentrations of drug, and at 72h TEER, Cilia beating frequency (CBF) and toxicity (LDH) were measured.
